# Variant Polycomb complexes in *Drosophila* consistent with ancient functional diversity

**DOI:** 10.1101/2022.04.29.490092

**Authors:** Hyuckjoon Kang, Janel R. Cabrera, Barry M. Zee, Heather A. Kang, Jenny Marie Jobe, Maeve B. Hegarty, Aurelie E. Barry, Alexander Glotov, Yuri B. Schwartz, Mitzi I. Kuroda

## Abstract

Polycomb group (PcG) mutants were first identified in *Drosophila* based on their failure to maintain proper *Hox* gene repression during development. The proteins encoded by the corresponding fly genes mainly assemble into one of two discrete Polycomb Repressive Complexes: PRC1 or PRC2. However, biochemical analyses in mammals have revealed alternative forms of PRC2, and multiple distinct types of non-canonical or variant PRC1. Through a series of proteomic analyses, we identify analogous PRC2 and variant PRC1 complexes in *Drosophila*, as well as a broader repertoire of interactions implicated in early development. Our data provide strong support for the ancient diversity of PcG complexes, and a framework for future analysis in a longstanding and versatile genetic system.

## Introduction

The Polycomb Group (PcG) regulatory complexes, PRC1 and PRC2, each encompass numerous alternative subunits and configurations in mammalian cells (Aranda et al., 2015; Gil and O’Loghlen, 2014; Holoch and Margueron, 2017; Piunti and Shilatifard, 2021; Turner and Bracken, 2013; van Mierlo et al., 2019). This has not been explored to the same extent in *Drosophila*, where Polycomb complexes comprise a reduced number of paralogous and accessory subunits (Kuroda et al., 2020). In the absence of extensive analyses, it has been assumed that PcG complexes have greatly diversified in mammals. However, numerous subunits of alternative PcG complexes are highly conserved, including the RING1 and YY1 Binding Protein (RYBP) and Polycomb Group RING Finger (PCGF) proteins, which have ancient origins (Gahan et al., 2020).

The previously described *Drosophila* dRAF and PhoRC complexes have multiple subunits in common with mammalian vPRC1.1 and vPRC1.6, respectively (Alfieri et al., 2013; Lagarou et al., 2008). However, several subunits thought to play defining roles in mammals were not detected in the fly complexes, including RYBP. Furthermore, orthologous PRC1.3/5 complexes have not been reported. These observations support the need for additional analyses in *Drosophila*.

Here we use crosslinking, tandem affinity purification, and mass spectrometry (BioTAP-XL) to discover that fly embryos utilize RYBP and three PCGF subunits, Psc (CG3886), Su(z)2 (CG3905), and l(3)73Ah (CG4195) to assemble complexes related to all previously described variant PRC1 subtypes. CG14073(BCOR) is a signature subunit of PRC1.1, and CG8677 (RSF1) is a newly identified interactor. Fly PRC1.3/5 may have a conserved role in the nervous system based on tay (CG9056), its defining subunit. Sfmbt (CG16975) interactions, which encompass the previously described PhoRC-L, suggest surprisingly broad modularity of a potential fly vPRC1.6.

We also confirm the modularity of PRC2 in *Drosophila*, with Pcl (CG5109) and Scm (CG9495) restricted to PRC2.1 and Jarid2 (CG3654) and jing (CG9397) restricted to PRC2.2. The conservation appears to extend to the association of PRC2.1 with stable repression, and PRC2.2 with a heterogenous or transitional role. Phenotypes from overexpression of jing or compensatory knockdown of Jarid2 provide further evidence for the importance of a proper balance between PRC2.1 and PRC2.2 during development.

## Results and Discussion

### RYBP and l(3)73Ah are core subunits of *Drosophila* vPRC1 complexes

Biochemical analyses have defined two major classes of PRC1 complexes in mammals. Canonical PRC1 complexes (cPRC1.2 and cPRC1.4) function in the maintenance of stable repression that is critical for the classical role of *Hox* gene repression. Non-canonical or variant PRC1 complexes (ncPRC1 or vPRC1) are responsible for the majority of ubiquitination of H2A on lysine 119 (H2AK119ub) and appear to have diverse functions, including both activation and repression in the case of PRC1.3/5 (Cohen et al., 2018; Fursova et al., 2019; Gao et al., 2014; Gao et al., 2012; Scelfo et al., 2019; Zhao et al., 2017). In mammals, either RING1A or RING1B is present in all canonical and variant PRC1 complexes. In contrast, RYBP or YAF2 (YY1 Associated Factor 2) is limited to variant PRC1 complexes (Wang et al., 2010), and six PCGF proteins define the two canonical and four variant PRC1 complexes (Gao et al., 2012) (**Fig. 1A**).

**Figure 1.**
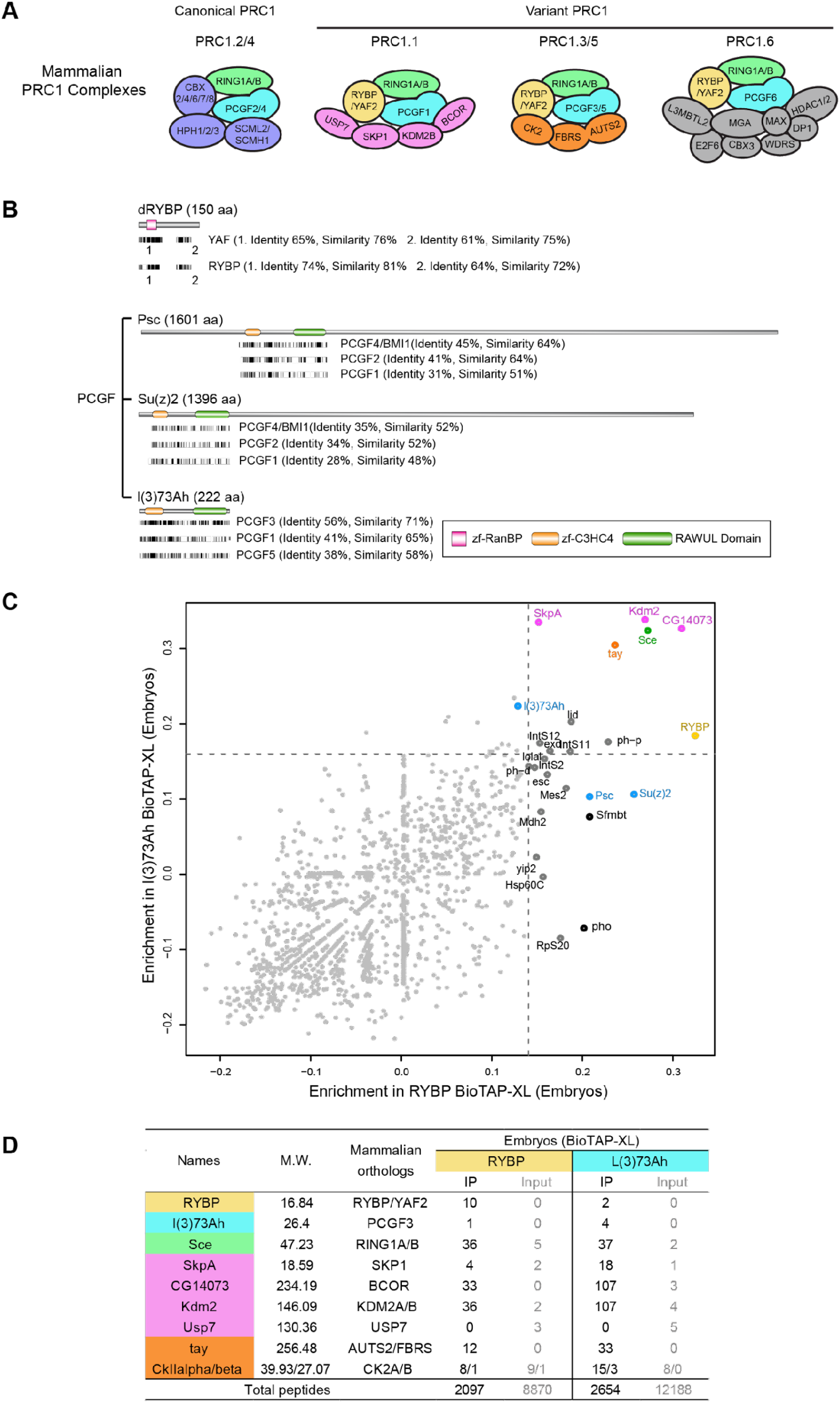
**(A)** Mammalian PRC1 complex components. PRC1 complexes contain core subunits RING1A/B and PCGF, and variant PRC1 (vPRC1) complexes commonly contain RYBP or YAF2. Each PRC1 subtype is defined by distinct PCGF proteins and additional accessory proteins, as indicated. **(B)** Top, sequence alignment of *Drosophila* RYBP (NP_001286742.1) with mammalian YAF2 (XP_011536030.1) and RYBP (NP_036366.3). Bottom, alignment of Psc (NP_001286368.1), Su(z)2 (NP_001260933.1), and l(3)73Ah (NP_001246797.1) with mammalian PCGF orthologs, PCGF1 (NP_116062.2), PCGF2 (NP_001356543.1), PCGF3 (NP_001304765.1), PCGF4/BMI1 (NP_005171.4), and PCGF5 (NP_001243478.1). Identical sequences of conserved regions are depicted as black lines, and percent sequence identity and percent similarity (identity + conservative substitutions) between two protein sequences are described in parentheses. Conserved domains are also shown. **(C)** Enrichment plots (log10 fold enrichment of normalized spectral abundance factors (NSAF) in pulldown with respect to input) for proteins (individual dots) identified in RYBP and l(3)73Ah BioTAP-XL experiments from *Drosophila* embryos. Dashed lines denote the 99th percentile threshold of enriched proteins in RYBP pull-down (x-axis) and in l(3)73Ah pull-down (y-axis). **(D)** Peptide counts of *Drosophila* PRC1.1 and PRC1.3 subunit orthologs co-purified by RYBP and l(3)73Ah embryonic BioTAP-XL IP compared to input. Proteins are color-coded according to their mammalian vPRC1 subunit orthologs in (A). Counts of all peptides detected in BioTAP-XL pull-downs and inputs are indicated as total peptides at the bottom of table.

As in mammals, Sce (CG5595, also known as dRING) is a conserved core subunit of *Drosophila* PRC1. In addition, *Drosophila* RYBP (CG12190) shows strong similarity to the mammalian paralogs, RYBP and YAF2 (**Fig. 1B**), and is known to interact with Sce (Fereres et al., 2014). The orthologous relationships of PCGF proteins in flies appear somewhat less straightforward. The two well-studied orthologs, Psc and Su(z)2, play central but partially redundant roles in cPRC1(Beuchle et al., 2001; Kassis et al., 2017; King et al., 2005; Li et al., 2009) and are most similar to mammalian PCGF4, PCGF2, and PCGF1 (**Fig. 1B and Fig.S1**). The most evolutionarily conserved fly PCGF protein, l(3)73Ah (Gahan et al., 2020; Lee et al., 2015) has not been studied with regard to its ability to participate in PcG complexes, but is most related to PCGF3, the most ancient of the PCGF family (Gahan et al., 2020). Finally, all fly PCGF orthologs are relatively distant from mammalian PCGF6 **(Fig.S1**)

Ideally, we would identify all candidate PRC1 subunits in flies by affinity purifying Sce-associated complexes. However, we were unable to recover functional Sce protein after tagging either the N or C terminus with the BioTAP epitope **(Fig. S2A)**. Therefore, we proceeded to tag RYBP and l(3)73Ah, established that they were functional as fusion proteins by viability rescue tests **(Fig. S2A)**, and performed BioTAP-XL affinity purification and mass spectrometry in the respective transgenic 12-24 hr embryos (**Fig. 1C**). Perhaps due to the small sizes of the two bait proteins, only 10 peptides of RYBP and 4 peptides of l(3)73Ah were recovered in their corresponding affinity purifications. Nevertheless, the pulldowns enriched for many candidate vPRC1 subunits at substantive levels (**Fig. 1D**). For example, several orthologs of mammalian PRC1.1 and PRC1.3/5 were co-purified in both embryonic RYBP and l(3)73Ah BioTAP-XL pull-downs, strongly suggesting that these variant complexes are conserved in *Drosophila*. The RYBP pull-down additionally co-purified Sfmbt, pho (CG17743) and Psc/Su(z)2 as well as PRC1.1. and PRC1.3/5 subunit orthologs, suggesting that RYBP could be a core subunit all vPRC1 complexes, as it is in mammals.

### Kdm2 and CG14073 (BCOR) are unique to vPRC1.1, while tay defines vPRC1.3/5

From our previous results, we could not exclude the possibility that orthologs of mammalian PRC1.1 and PRC1.3/5 subunits enriched in both RYBP and l(3)73Ah pull-downs might be subunits of one composite complex in *Drosophila*. Therefore, we performed Kdm2 (CG11033) BioTAP-XL in embryos to determine whether its presence could discriminate between fly vPRC1.1 and PRC1.3/5. We were unable to test BioTAP-tagged Kdm2 for functionality, as Kdm2 mutants are viable (Cohen et al., 2018); however, we successfully co-purified Kdm2-interacting orthologs of mammalian PRC1.1. Whether interactions are expressed as enrichment plots (**Fig. 2A and 2B**) or in a heat map (**Fig. 2C**) it is apparent that tay, the fly ortholog of mammalian signature PRC1.3/5 subunits AUTS2/FBRS/FBRSL1, was not co-purified in BioTAP-XL pull-down of Kdm2. From this we conclude that PRC1.1 and PRC1.3/5 exist as separate vPRC1 complexes in *Drosophila*, as in mammals (**Fig. 2D**). Interestingly, fly *tay* was discovered in a behavioral screen of locomotor mutants (Poeck et al., 2008), while AUTS2 is named for its association with autism in humans (Gao et al., 2014; Oksenberg and Ahituv, 2013), suggesting a conserved link to the nervous system in the two evolutionarily distant species.

**Figure 2.**
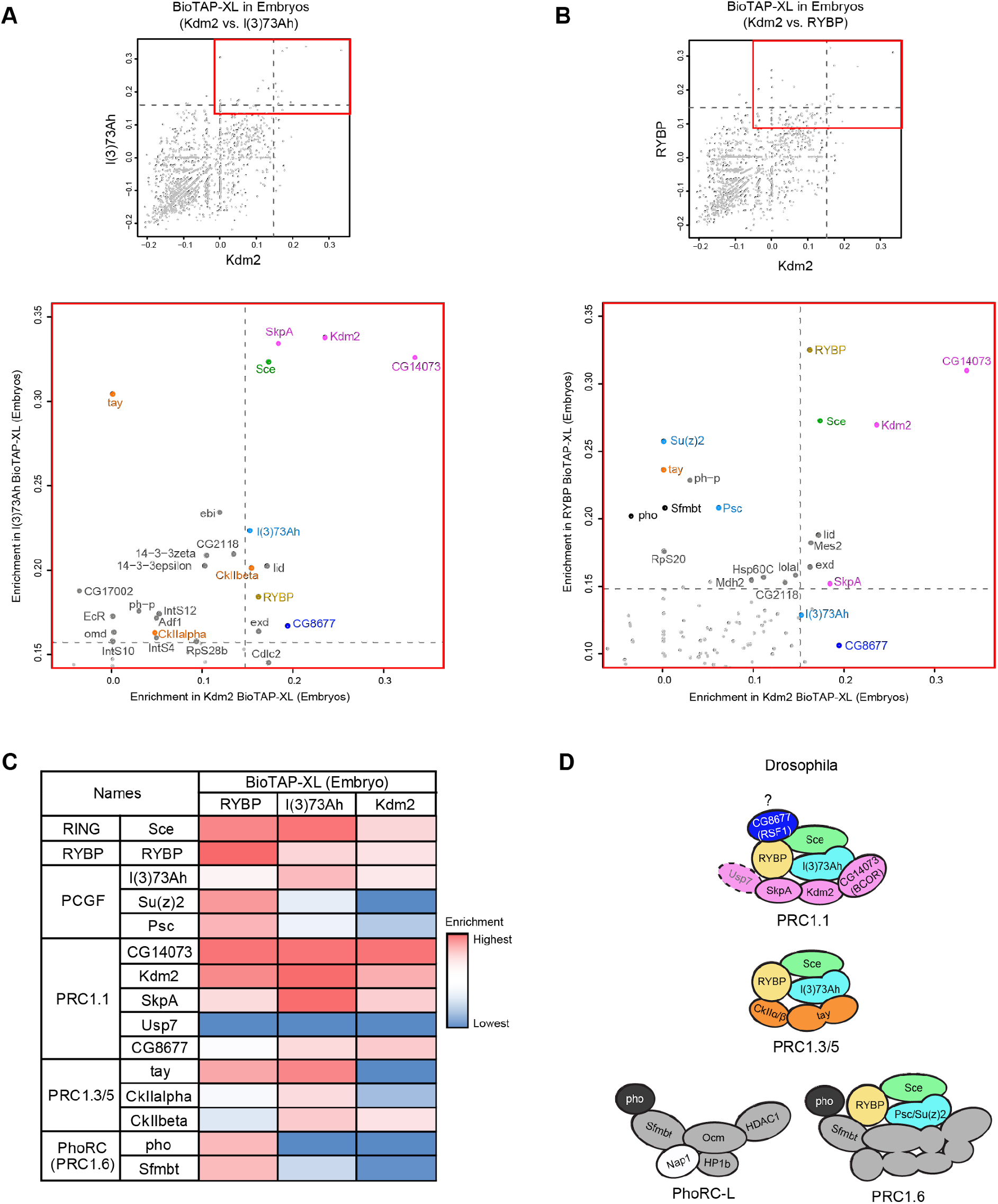
**(A-B)** Scatterplot comparing enriched proteins from Kdm2 and l(3)73Ah BioTAP-XL from 12-24hr embryos **(A)**. Scatterplot comparing enriched proteins from Kdm2 and RYBP pulldowns from 12-24hr embryos **(B)**. Dashed lines on x-axis denote 99th percentile threshold of enriched proteins in Kdm2 pulldown (x-axis) and dashed lines on y-axis denote 99th percentile threshold of enriched proteins by l(3)73Ah and RYBP in (A) and (B), respectively. Coordinates represent log10 fold enrichment of normalized spectral abundance of proteins in BioTAP-XL affinity purification compared to input. Red boxes indicate areas of scatterplots that are enlarged in bottom panels and protein names are color-coded according to the vPRC1 subunit color scheme in Fig. 1A and Fig. 2D. **(C)** Heat map of enrichment of selected vPRC1 subunits co-purified from RYBP, l(3)73Ah and Kdm2 BioTAP-XL. For relative abundance comparison, the NSAF enrichment value of each protein is divided by the NSAF enrichment value of CG14073 (BCOR) which is the top hit common to all three pulldowns. **(D)** Illustration of *Drosophila* vPRC1 complexes from BioTAP-XL mass spec analysis of the three bait proteins RYBP, l(3)73Ah, and Kdm2 organized similarly to their mammalian complexes in Fig. 1A. Usp7 (not copurified by three bait proteins) is depicted by a dashed line. CG8677, indicated by a question mark, is a potential *Drosophila*-specific subunit of PRC1.1.

Although the previously described dRAF complex was identified after Kdm2 co-purification with Psc (Lagarou et al., 2008), our Kdm2 and l(3)73Ah BioTAP-XL pull-downs did not enrich for Psc or Su(z)2, while orthologs of PRC1.1 subunits, eg. CG14073 (BCOR) and SkpA (CG16983), strongly interacted with Kdm2. We cannot exclude the existence of the dRAF complex in flies, but Kdm2 seems to mainly form a complex with CG14073, SkpA, Sce and l(3)73Ah in *Drosophila* embryos.

Unexpectedly, the Kdm2 BioTAP-XL pulldown also strongly co-purified CG8677, the *Drosophila* ortholog of RSF1(Remodeling and spacing factor 1). In mammals, RSF1 specifically recognizes H2AK119ub, the chromatin mark catalyzed principally by vPRC1 (Zhang et al., 2017). As l(3)73Ah also co-purified CG8677, this result supports the discovery of RSF1 as a PcG related H2Aub reader (Zhang et al., 2017). Surprisingly, Usp7 (CG1490), a ubiquitin-specific protease, was absent from our purifications, raising the possibility that the recovery of CG8677 (RSF1) could be related to stabilization or accessibility of H2Aub.

### Potential relationship of PhoRC to PRC1.6, with an expanded repertoire of interactions

Based on the prior discovery of PhoRC and PhoRC-L as stable, soluble Sfmbt complexes (Alfieri et al., 2013; Klymenko et al., 2006) (**Fig. 2D**), and on recovery of pho, Sfmbt, Psc, and Su(z)2 in our RYBP pulldown (**Fig. 2B and 2C**), we created BioTAP-Sfmbt as a candidate bait to further explore the possible relationship between PhoRC-L and vPRC1.6 in flies. We found that BioTAP-Sfmbt associated with larval polytene chromosomes as expected for a PcG protein **(Fig. S2B)** but failed to rescue mutants to viability **(Fig. S2A)**. Previously, a similarly tagged Sfmbt protein also failed to rescue mutant animals, but restored *Hox* gene repression in mutant clones and was used successfully to affinity-purify PhoRC-L (Alfieri et al., 2013). Therefore, we proceeded with proteomic analysis of BioTAP-tagged Sfmbt in both embryos and a stable S2 cell line.

Unexpectedly, the Sfmbt pulldowns generated a much longer list of interactors than typical for the relatively discrete PcG complexes we have analyzed in the past. Subunits of fly PhoRC-L and orthologs of mammalian PRC1.6 subunits were enriched, but broadly distributed in the enrichment rankings (**Fig. 3A and 3B**). To confirm this, we compared the enrichments between the embryo and S2 cell experiments **(Fig. 3C)**. Dashed lines denote the top 1% enrichment in S2 cells (x-axis) and in 12-24hr embryos (y-axis) showing general agreement between the two distinct biological samples. The total peptide counts of selected proteins enriched in Sfmbt BioTAP-XL pulldowns are shown in **Fig. 3D**, compared to their cognate inputs. Two notable orthologs of mammalian vPRC1.6 (**Fig. 1A**) were missing from our Sfmbt pulldown: RYBP and MAX. We previously found that when RYBP was used as the BioTAP-XL bait, it was a strong interactor of Pho and Sfmbt, (**Fig. 2C**) even when RYBP itself was represented by only 10 peptides (**Fig.1D**). Therefore, RYBP might not be identified by the reciprocal Sfmbt pulldown due to its small size and thus fewer peptides for detection after crosslinking. *Drosophila* Max (CG9648) might also be undetected due to its small size (18.5 kDa). However, it seems likely that Max is not a subunit of *Drosophila* Sfmbt complexes, because the bHLHZip domain for heterodimerization between MGA and MAX is absent in *Drosophila* MGA orthologue ocm (CG3363) (Hurlin et al., 1999).

**Figure 3.**
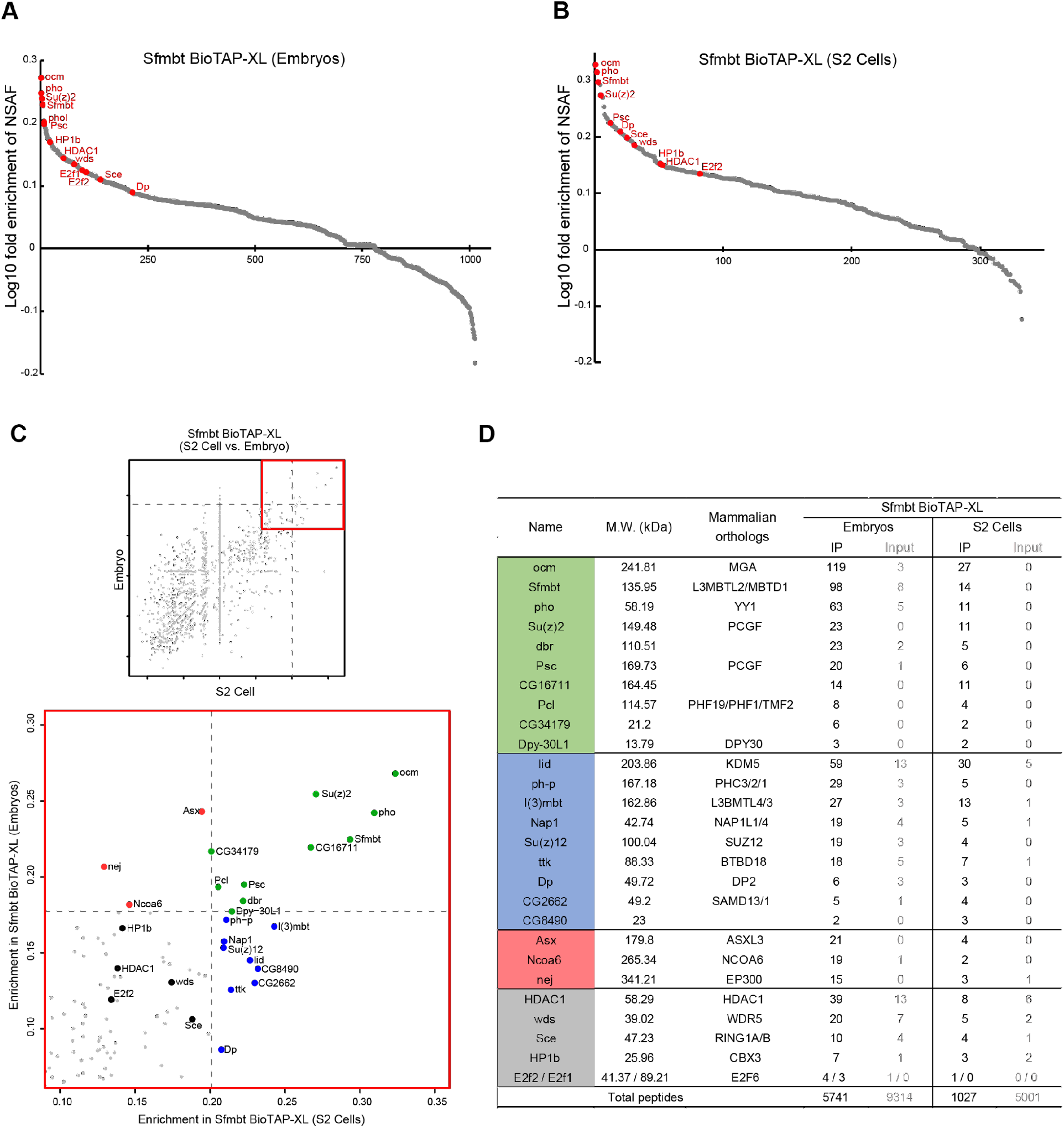
**(A - B)** Log10 fold enrichment of relative abundance for proteins identified in Sfmbt BioTAP-XL experiments from 12-24hr embryos **(A)** and from S2 cells **(B)**. Orthologs of mammalian PRC1.6 subunits as well as PhoRC subunits are highlighted in red. **(C)** Scatter plot of Sfmbt pull-down enrichments from embryos and S2 cells normalized to their inputs. Dashed lines represent the 99th percentile of proteins enriched by Sfmbt BioTAP-XL in embryos (x-axis) and in S2 cells (y-axis). Red box indicates the area of the scatterplot enlarged in the bottom panel. Top 1% of interactors in both embryos and S2 cells are highlighted in green. Protein lists enriched in either S2 cells or embryos (top 1%) are highlighted in blue and red, respectively. Orthologs of mammalian PRC1.6 subunits which are not top 1% interactors but are still enriched are highlighted in black. **(D)** Total peptide counts of select proteins enriched from Sfmbt BioTAP-XL experiment in embryos and in S2 cells compared to peptide counts from respective inputs. Protein names are highlighted with different colors according to colored dots in scatter plot (C). Total peptide counts from pull-downs are indicated at the bottom of table.

In addition to the orthologs previously implicated in mammalian vPRC1.6, we found *Drosophila-specific* interactors (dbr (CG11371), CG16711, CG34179) and proteins with Sterile alpha motif (SAM) domains (l(3)mbt (CG5954), CG2662, ph-p). Fly Sfmbt itself has a SAM domain, which is absent from its mammalian L3MBTL2 counterpart (**Fig. 4A**). Thus, it may not be surprising that Sfmbt can interact with other SAM domain proteins through homo- or heteromeric interaction, a known property of SAM domains (Kim and Bowie, 2003). In fact, Sfmbt was one of the most enriched proteins in our previous analysis of BioTAP-XL Scm (Kang et al., 2015) and Scm mediates a trimeric complex with Sfmbt and ph-p through its SAM domain (Frey et al., 2016). Consistent with these key interactions, our Sfmbt pulldown also enriched for subunits of cPRC1 (ph-p (CG18412)) and PRC2 (Pcl (CG5109), Su(z)12 (CG8013)), and previous pulldowns of cPRC1 and PRC2 identified Sfmbt and pho (Kang et al., 2015; Strubbe et al., 2011). Taken together, our results are consistent with the ability of PhoRC to serve as an anchor for PcG complexes at PREs (Frey et al., 2016).

**Figure 4.**
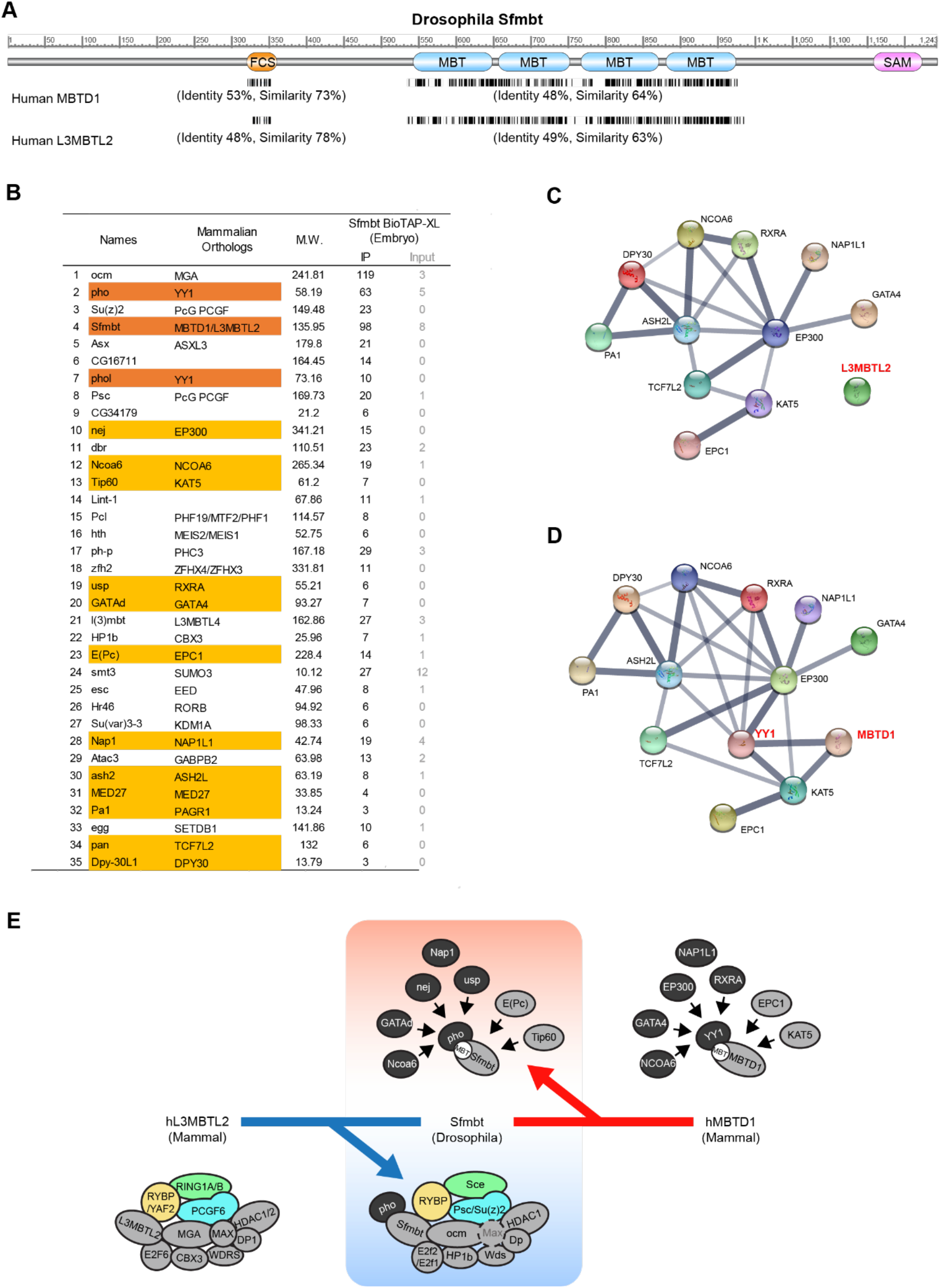
**(A)** Conserved domains in *Drosophila* Sfmbt and sequence alignments with human MBT domain orthologs. Sequence identities between *Drosophila* Sfmbt protein sequence (NP_001137821.1) and two human orthologs, MBTD1 (XP_011523224.1) and L3MBTL2 (NP_001137821.1) are shown as black lines. Percent sequence identity and percent similarity in domains are also described in parentheses. **(B)** Total peptide counts of top 35 proteins enriched from Sfmbt BioTAP-XL in embryos. Sfmbt and pho/phol proteins are highlighted in orange. Among non-PcG proteins, proteins with mammalian orthologs that have known interactions are highlighted in yellow. **(C)** STRING protein-protein interaction network for human orthologs of yellow-highlighted proteins in (B). Only known interactions (from curated databases and experimentally determined) are used as interaction sources, and the thickness of the gray lines indicates strength of supporting data. L3MBTL2 (human ortholog of Sfmbt) is not connected with the interaction network group. **(D)** Combined interaction network of MBTD1 (human ortholog of Sfmbt) and YY1 (human ortholog of pho/phol). **(E)** *Drosophila* Sfmbt may have properties of both L3MBTL2 and MBTD1 orthologs and may be able to interact with binding partners of each ortholog.

### Sfmbt complexes are linked to co-activators in embryos

Interestingly, we found interactions with co-activators and additional chromatin regulators in embryos that were diminished or absent in S2 cells, suggesting that Sfmbt complexes may have functions beyond PcG repression during development (**Fig. 3D, 4B and Fig. S3**). Furthermore, while Sfmbt is a *Drosophila* ortholog of L3MBTL2, it is also equidistant in identity and similarity to mammalian MBTD1 (**Fig. 4A**). When we examined the top 35 proteins enriched by Sfmbt pulldown specifically in embryos, we noted that those highlighted in yellow (**Fig. 4B**) map to a common STRING protein-protein interaction network in mammals (**Fig. 4C**, and **4D**). The interaction network is not connected to L3MBTL2 **(Fig. 4C)**. However, YY1 provides a potential bridge between the network and MBTD1 **(Fig. 4D)**, as it can bind to a helical groove of MBTD1 in vitro, although with a relatively weak binding affinity (Alfieri et al., 2013). Taken together, we can speculate that Sfmbt may be the *Drosophila* ortholog of both L3MBTL2 and MBTD1. Interestingly, the MBTD1 network and our pulldown results include several prominent interactors implicated in activation. These include nej (CG15319), the ortholog of the EP300/CBP acetyltransferases, and Tip60 (CG6121) and E(Pc) (CG7776), orthologs of members of the NuA4/TIP60 acetyltransferase complex. That pho/phol could be linked to transcriptional activation would be consistent with their presence on PRE regulatory sequences in both repressive and activating contexts (De et al., 2019; Fujioka et al., 2008; Ghotbi et al., 2021; Kahn et al., 2014; Kassis and Brown, 2013; Loubiere et al., 2020; Schuettengruber et al., 2009). Furthermore, the pho/phol mammalian ortholog, YY1 is named for its dual function in the two opposing gene regulatory states (Shi et al., 1991).

In summary, we provide strong evidence that *Drosophila* has counterparts for all variant PRC1 complexes found in mammals, with most subunits conserved. Notably, PCGF proteins don’t have a strict one to one correspondence between flies and mammals, with individual subtypes participating in multiple configurations in *Drosophila*. Furthermore, when using Sfmbt as the bait protein in affinity purifications, we detected many additional interactors that could be related to Sfmbt playing the roles of both mammalian L3MBTL2 and MBTD1, and as an important PcG anchor at PREs. We favor a model in which a hypothetical fly vPRC1.6 is not a single entity, but instead encompasses modules with distinct functions which will require additional molecular analysis and dissection in the future **(Fig. 4E)**.

### Conservation of modular PRC2.1 and PRC2.2 complexes

Mammals express a core PRC2 complex, with two main alternatives incorporating mutually exclusive accessory proteins (Holoch and Margueron, 2017; van Mierlo et al., 2019). PRC2.1 and PRC2.2 appear redundant in mammalian ESCs, but their functions diverge during differentiation (Healy et al., 2019; Petracovici and Bonasio, 2021). The presence of mutually exclusive PRC2.1 and PRC2.2 accessory subunits in *Drosophila* has been inferred from several previous studies (Herz et al., 2012; Kalb et al., 2014; Nekrasov et al., 2007). Our previous BioTAP-XL analysis of *Drosophila* E(z) also recovered the four PRC2 core subunits as well as Scm, Pcl, Jarid2, and jing (Kang et al., 2015). The latter three proteins are the orthologs of mammalian accessory subunits PHF1/PHF19/MTF2, JARID2, and AEBP2. To confirm whether these accessory subunits form orthologous PRC2.1 and PRC2.2 complexes in *Drosophila*, we prepared transgenic fly lines expressing BioTAP tagged Pcl, Jarid2, and jing. The tagged transgenes rescued their corresponding homozygous lethal mutants to viability, demonstrating that they each expressed functional proteins (**Fig S2A**), and Pcl and Jarid2 enriched for PRC2 complexes as expected (**Fig. 5A**). However, pulldown of epitope tagged jing failed to recover even the bait protein (data not shown). Interestingly, Pcl and Scm co-purified each other but not Jarid2 and jing, and Jarid2 only co-purified jing but not Pcl and Scm (**Fig. 5A**). The graphed comparison of Pcl and Jarid2 BioTAP-XL enrichments reveal the common core subunits on the diagonal and the mutually exclusive accessory subunits as top 1% interacting partners in their respective pulldowns (**Fig. 5B**).

**Figure 5.**
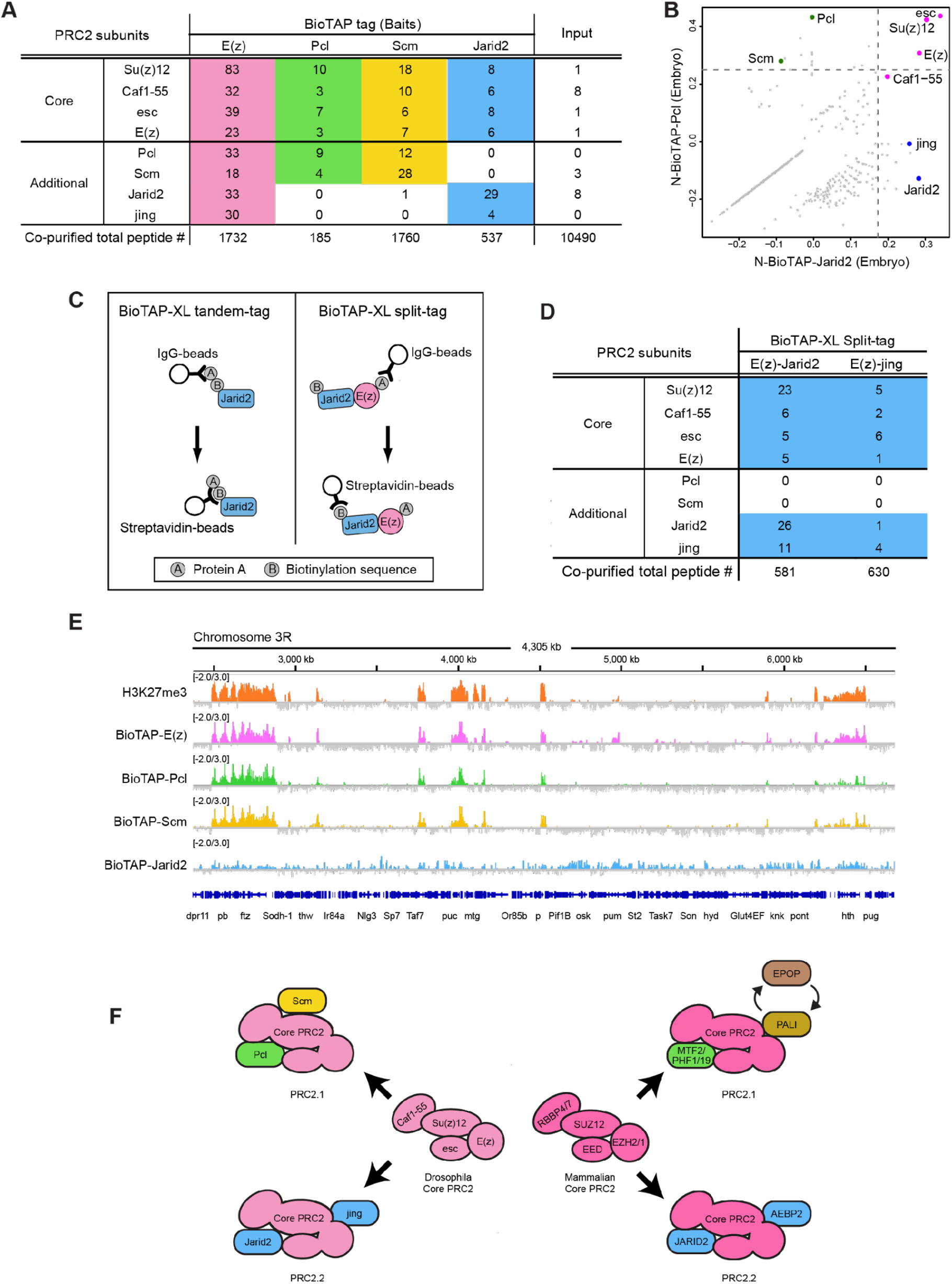
**(A)** Peptide counts of PRC2 subunits recovered from BioTAP-XL pull-downs of bait proteins in embryos compared with peptide counts of input. Total peptides are indicated at the bottom of table. **(B)** Scatter plot showing Jarid2 and Pcl pull-down enrichment (normalized to total embryonic chromatin input). PRC2 core subunits are highlighted in pink, and additional subunits of PRC2.1, and PRC2.2 are highlighted in green and blue, respectively. **(C)** Schematic of co-purification strategies of BioTAP-XL tandem-tag and split-tag. **(D)** Peptide counts of PRC2 subunits co-purified by BioTAP-XL split tag pulldowns. **(E)** Genome browser view of a region of chromosome 3R showing co-localization of Pcl and Scm with E(z) and H3K27me3 in embryos, while Jarid2 is widely dispersed across the genome. **(F)** Cartoon showing conservation of PRC2 complexes between *Drosophila* and mammals. Scm and PALI are not conserved orthologs but may have functional similarity via their separate interaction with G9A methyltransferase.

Although BioTAP-XL using jing as the bait was unsuccessful, we knew that jing was clearly detected in our original E(z) pulldown (**Fig. 5A**). Therefore, we designed a sequential purification approach to further validate the existence of PRC2.2 in *Drosophila*. We split the dual BioTAP-tag between two bait proteins **(Fig. 5C**), tagging E(z) with Protein A, and either Jarid2 or jing with the biotinylation target sequence. The sequential tandem tag purifications of either ProteinA-E(z) /Bio-Jarid2 or ProteinA-E(z)/ Bio-jing resulted in relatively low numbers of total peptides, but confirmed Jarid2 and jing as genuine PRC2.2 subunits which were mutually exclusive with Pcl and Scm (**Fig.5D**).

To investigate whether the conservation of PRC2.1 and 2.2 in mammals and flies extends to divergent genomic occupancy and potentially function during differentiation, we mapped the genomic occupancy of BioTAP-Pcl and -Jarid2 in 12-24 hour embryos. Consistent with foundational work in mammalian cells (Petracovici and Bonasio, 2021), occupancy of *Drosophila* BioTAP-Pcl (PRC2.1) correlates with stable repression and H3K27me3. In contrast, occupancy of BioTAP-Jarid2 (PRC2.2) suggests a heterogenous or transitional role (**Fig. 5E and Fig.S4D-F)**. The genetically functional BioTAP-Jarid2 binding pattern is surprisingly distinct from PRC2.1 and H3K27me3; however we validated it by comparison to the binding pattern of functional Jarid2-GFP from the ENCODE project (**Fig. S4A-C)**. These surprising results could be consistent with a transitional role for PRC2.2 in bringing PRC2 methyltransferase activity to transcriptionally active regions (Kasinath et al., 2021; Petracovici and Bonasio, 2021).

Based on the importance of a balance between PRC2.1 and PRC2.2 in mammalian cells (Youmans et al., 2021), we asked whether overexpressing jing or Jarid would affect normal development. We found that overexpression of UAS-Jarid driven by engrailed-GAL4 was viable with no detectable phenotype, but that overexpression of UAS-jing yielded no adult progeny (**Fig. S4G**). Interestingly, Jarid2 partial RNAi knockdown suppressed the lethality caused by overexpression of jing (**Fig. S4G**), consistent with PRC2.2 as a functional unit whose proper levels are critical during development.

As shown in **Fig. 5F** our mass spec data demonstrate similar compositions of PRC2.1 and PRC2.2 in flies and mammals, with the notable substitution of vertebrate-specific PALI1 or EPOP for Scm in *Drosophila*. While Pcl, Jarid2, and jing correspond to clearly orthologous PRC2 subunits in mammals, Scm may play a functionally conserved role without sequence homology, as PALI1 and Scm both have strong associations with orthologous G9A methyltransferases (Alekseyenko et al., 2014; Conway et al., 2018; Kang et al., 2015). Furthermore, SCML2, a potential mammalian ortholog of Scm, is known to regulate PRC2 activity during spermatogenesis, suggesting that a common primordial interaction may occur in the germline (Maezawa et al., 2018).

## Concluding Remarks

In summary, our data provide strong support for an ancient functional diversity of PcG complexes, with remarkable conservation between mammals and *Drosophila*. Historically, this was not evident as many of the cognate mutants failed to display the classical homeotic phenotype first associated with defects in Polycomb repression in *Drosophila*. In the future, it will be interesting to dissect the germline functions and potential pleiotropy of these undoubtedly important regulators of cellular identity during development.

## Acknowledgments

We thank R. Tomaino for expert help with protein samples (Taplin Mass Spectrometry Facility, Harvard Medical School), and J. Müller, HM. Herz, W. Bender, J. Culi, and D. Montell for generous sharing cDNA and/or fly stocks. We are grateful to J. Kassis for fly stocks and numerous helpful discussions. We thank the TRiP at Harvard Medical School (National Institutes of Health/National Institute of General Medical Sciences R01-GM084947) and the BDSC at Indiana University (National Institutes of Health P40OD018537) for providing fly stocks used in this study. This work was supported by an NIH grant to MIK (R35-GM126944) and a Swedish Research Council grant to YBS (2021-04435).

## Material and methods

### Fly Genetics

#### Transgenic fly lines

The pFly vector was used as the backbone to construct all transgenes (Wang et al., 2013). For expression of Sce, RYBP, Sfmbt, Kdm2, Pcl, E(Z), and jing, cDNA fragments were cloned under control of the α-tubulin 1 promoter. Jarid2 and l(3)73Ah were prepared from a ~13 kb genomic fragment amplified from BAC CH321-48A06 and a ~3.7 kb region amplified from *Drosophila* genomic DNA, respectively. The genomic fragments encompassed the promoter, coding, upstream and downstream regions. In most cases, we prepared both N-terminal and C-terminal BioTAP (ProteinA + Biotinylation target sequences) transgenic lines for each bait protein. Neither N- or C-BioTAP Sce could rescue mutant lethality. N-BioTAP tagged RYBP, Sfmbt, Jarid2 and Pcl and C-BioTAP tagged l(3)73Ah and Kdm2 were used for BioTAP-XL affinity purification in this study. For split tag experiments, N-terminal Protein A was added to an E(z) transgene (N-ProteinA-E(z)), and biotinylation target sequences were added to the N-terminus of either jing or Jarid2 transgenes (N-Biotin-jing and N-Biotin-Jarid2). Injections of all transgenes were performed at Bestgene Inc (https://www.thebestgene.com/), with the transgenes integrated into site-specific attP-docking sites by PhiC31 integrase-mediated transgenesis systems (N-BioTAP-RYBP, N-BioTAP-Sfmbt, N-BioTAP-Pcl and N-Biotin-jing transgenes: BDSC #9732 [76A2], N- and C-BioTAP-Sce, C-BioTAP-l(3)73Ah, C-BioTAP-Kdm2, N-BioTAP-Jarid2 and N-ProteinA-E(z): BDSC#9736 [53B2], and N-Biotin-Jarid2; BDSC#9748 [62E1]). For split tag purification, transgenic fly lines stably-expressing both PoteinA-E(z) and Biotin-Jarid2 (or Biotin-jing) were made by crossing the N-ProteinA-E(z) transgenic fly line to N-Biotin-Jarid2 or N-Biotin-jing fly lines, followed by establishment of homozygous stocks.

#### Rescue tests and mutant strains

The viability rescue tests of BioTAP transgenics were described in Supplementary Figure S2A. Viability was assessed by the absence of balancer markers in adult flies expressing the mini-white marker linked to the transgenes. Mutant fly stocks were kindly provided by Dr. J. Muller (*Sfmbt^1^* & Df(2L)BSC30), Dr. HM Herz (*Jarid2^e03131^* & *Jarid2^MB00996^*), Dr. W.W. Bender (In(2R)Pcl^11^, Df(2R)Pcl7B, & *Pcl^R5^*), and Dr. D.J. Montell (*Jing^47H6^* & *Jing^22F3^*). Other fly stocks were obtained from the BDSC at Indiana University (*Sce^1^* #24618, Df(3R)IR16 #9529, *RYBP^KG08693^* #14968, Df(2R)BSC598 #25431, Df(2R)BSC326 #24351 and Jarid2-GFP #66754). Two new l(3)73Ah null alleles were generated using CRISPR/Cas9-mediated genome editing strategy described in (Kondo and Ueda, 2013). Briefly, the y1, w67c23; pWattB-L3-Dual-CRISPR flies, which had a transgene expressing two sgRNAs (L(3)73Ah-gRNA1: 5’-AUGCUCGCGCACGCUUGUACGUUUUAGAGCUAGAAAUAGCAAGUUAAAAUAAGGCUAGUCCGUUAUCAACUUGAAAAAGUGGCACCGAGUCGGUGCUUUUU -3’ and L(3)73Ah-gRNA2: 5’CGUGACGCGCUGGCGCUUCCGUUUUAGAGCUAGAAAUAGCAAGUUAAAAUAAGGCUAGUCCGUUAUCAACUUGAAAAAGUGGCACCGAGUCGGUGCUUUUU -3’) from ubiquitous U6 promoter integrated at M{3xP3-RFP.attP}ZH-51D landing site, were crossed to y2,cho2,v1; attP40{nos-Cas9}/CyO strain. Resulting attP40{nos-Cas9}/pWattB-L3-Dual-CRISPR; +/+ females were crossed to w1; If/CyO; MKRS/TM6,Tb and the progeny individually screened for editing events by PCR with L(3)73Ah-upstr1 (5’-CGCGACTAGTAGCAGGTACG -3’) and L(3)73Ah-dwnstr1 (5’-AAATGCAGCAAAGAAGGCGG -3’) primers which amplify 1479bp fragment from unedited chromosomes and 427bp fragment from chromosomes with expected precise deletion. Two null alleles 1(3)73Ah612 (deletion breakpoints: chr3L: 16588680-16589732) and l(3)73Ah1012 (deletion breakpoints: chr3L: 16588681-16589549) that remove the start codon and almost the entire open reading frame were selected for further experiments.

#### Overexpression and RNAi knockdown

The UAS-jing FL transgenic fly line used for overexpression of jing was the generous gift from Dr. J. Culi. Transgenic RNAi Project (TRiP) HMS02022 line (BDSC #40855) and P{Mae-UAS.6.11} Jarid2^LA00681^ line (BDSC # 22187) were used for Jarid2 knockdown and overexpression, respectively. These misexpression or RNAi knockdown lines were crossed with the engrailed (en)-GAL4 flies (gift of Dr. N. Perrimon).

### Polytene chromosome immunostaining

Salivary gland polytene chromosomes from third instar larvae were prepared by first fixation for 1min with 1% Triton X-100, 4% formaldehyde in PBS, followed by second fixation for 2 min with 50% acetic acid, 4% formaldehyde prior to squashing to spread the polytene chromosomes. Rabbit peroxidase antiperoxidase (PAP) antibody (1:100 dilution, Sigma) are used for detection of BioTAP-Sfmbt, and donkey anti-rabbit Alexa fluor 594 (1:500) were used as secondary antibody.

### BioTAP-XL

The BioTAP-XL protocol was performed as described in (Alekseyenko et al., 2015). Briefly, 12 to 24 hr old embryos of BioTAP and split-tag transgenic fly lines were collected and stored for up to 3 days at 4°C. Embryonic nuclei were cross-linked with 3% formaldehyde, snap-frozen in liquid nitrogen, and then stored at −80°C. These steps were repeated until nuclear extracts from ~40 g embryos were pooled. After sonication of extracts, the first purification step of interaction between IgG-agarose beads and Protein A was followed by the second step binding between Streptativdin-conjugated beads and biotin to purify the tagged bait proteins along with their protein interaction partners and associated genomic DNA. Bound protein complexes were trypsinized on bead and peptides were desalted with C18-STAGE tips (3M) as described previously (Zee et al., 2016) for liquid chromatography mass spectrometry (LC-MS) analysis. Most LC-MS files were searched using the SEQUEST algorithm with precursor mass tolerance of 20 ppm and fragment ion tolerance of 0.9 Da. Peptide identifications were filtered with XCorr ≥ 2 for z = 2 and ΔCorr ≥ 0.1. In the case of l(3)73Ah pull-down and its input, we used 0.03 Da for a fragment ion tolerance. Genomic localization of BioTAP-Pcl and -Jarid was determined by high-throughput DNA sequencing of libraries generated from BioTAP-XL purified genomic DNA using the NEBNext Ultra II DNA Library Prep Kit for Illumina (New England Biolabs, catalog no. E7645) and NEBNext Multiplex Oligos for Illumina (New England Biolabs, catalog no. E7335). The library samples were sequenced using an Illumina HiSeq 2500. Proteomic data sets have been deposited at Harvard Dataverse (https://dataverse.harvard.edu/dataverse/vPRC) and will be available in the PRIDE database upon publication.

### Proteomic analysis

To identify the proteins enriched by the BioTAP-tagged fusion proteins, a modification of the normalized spectral abundance factor (NSAF) was used to calculate enrichment ratios (pull-down/input) for each identified protein (Zybailov et al., 2006). The total number of identified peptides for a given protein were divided by the protein molecular weight (kilodalton) to control for protein size. To allow the calculation for proteins which were not recovered in the input sample, a pseudocount of 0.5 peptides was substituted for zero peptides. Then, the weight-normalized counts for each protein were divided by the sum of weight-normalized counts across all proteins in the sample. After natural log transformation of the value, the immunoprecipitation pulldown values were divided by the input values, log10-normalized, and multiplied by −1 to yield enrichments of interactors for the BioTAP-bait proteins

### ChIP-seq Analysis

ChIP-seq data for BioTAP-Jarid2 and BioTAP-Pcl have been submitted to the NCBI Gene Expression Omnibus repository under the accession number GSE201842. ChIP-seq data for E(z) and Scm were generated in our previous study (Kang et al., 2015); raw sequencing files are deposited in GEO (accession number GSE66183). Sequencing files for H3K37me3 were obtained from GEO (GSE47230) and those for Jarid2-GFP were obtained from http://encodeproject.org (Accession #s: ENCFF090YBK, ENCFF064HTW, ENCFF649NYX, ENCSR388YUZ).

Raw fastq files were preprocessed with Trimmomatic (Bolger et al., 2014) to remove Illumina adapters and quality filtered using the fastq_quality_filter function from FASTX-Toolkit (http://hannonlab.cshl.edu/fastx_toolkit) to retain reads with a quality score of ≥20 for at least 80% of bases. Reads were then aligned to the Drosophila genome (dm3) using Bowtie2 (Langmead and Salzberg, 2012) with default parameters. SAM files were processed to BAM files and reads with mapping quality (MAPQ) ≥20 were retained using Samtools (Li et al., 2009). BAM files were read normalized by RPKM (reads per kilobase per million mapped reads) and log2 ChIP/Input ratio binding profiles for all ChIP-seq replicates were generated using deepTools (Ramirez et al., 2016) bamCompare function for visualization using Integrative Genome Viewer (IGV) (Robinson et al., 2011). Correlation between biological replicates was assessed with deepTools multiBigwigSummary bins and plotCorrelation functions to compute Pearson correlation coefficient. Peaks were called using MACS2 (Zhang et al., 2008) for each experimental sample with matched input as control and a significance cutoff of -q 0.05. Overlapping peak regions were determined using the BEDTools software suite (Quinlan and Hall, 2010) and Venn diagrams were generated using the VennDiagram R package (Chen and Boutros, 2011).

---

**Figure S1.**
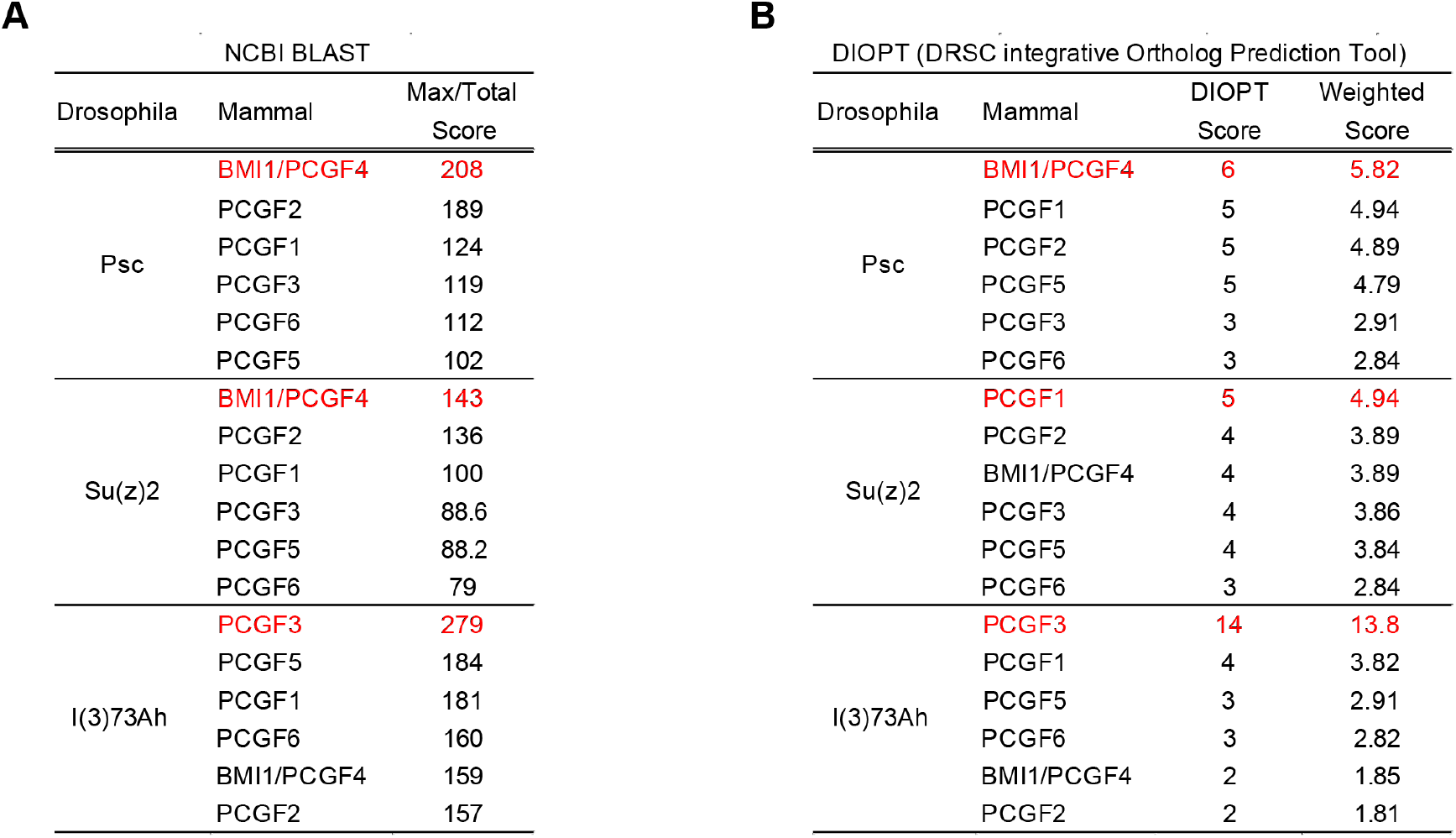
**(A and B)** *Drosophila* PCGF proteins and ranking of mammalian orthologs according to alignment score. While PCGF1, PCGF2, and BMI1/PCGF4 are in the top3 for Psc and Su(z)2 and PCGF1, PCGF3, and PCGF5 are in the top3 for l(3)73Ah, PCGF6 is relatively distant to all Drosophila PCGF proteins **(A)** Rankings from Max/total score of NCBI BLAST **(B)** Rankings from DIOPT score and Weighted Score of DRSC DIOPT.

**Figure S2.**
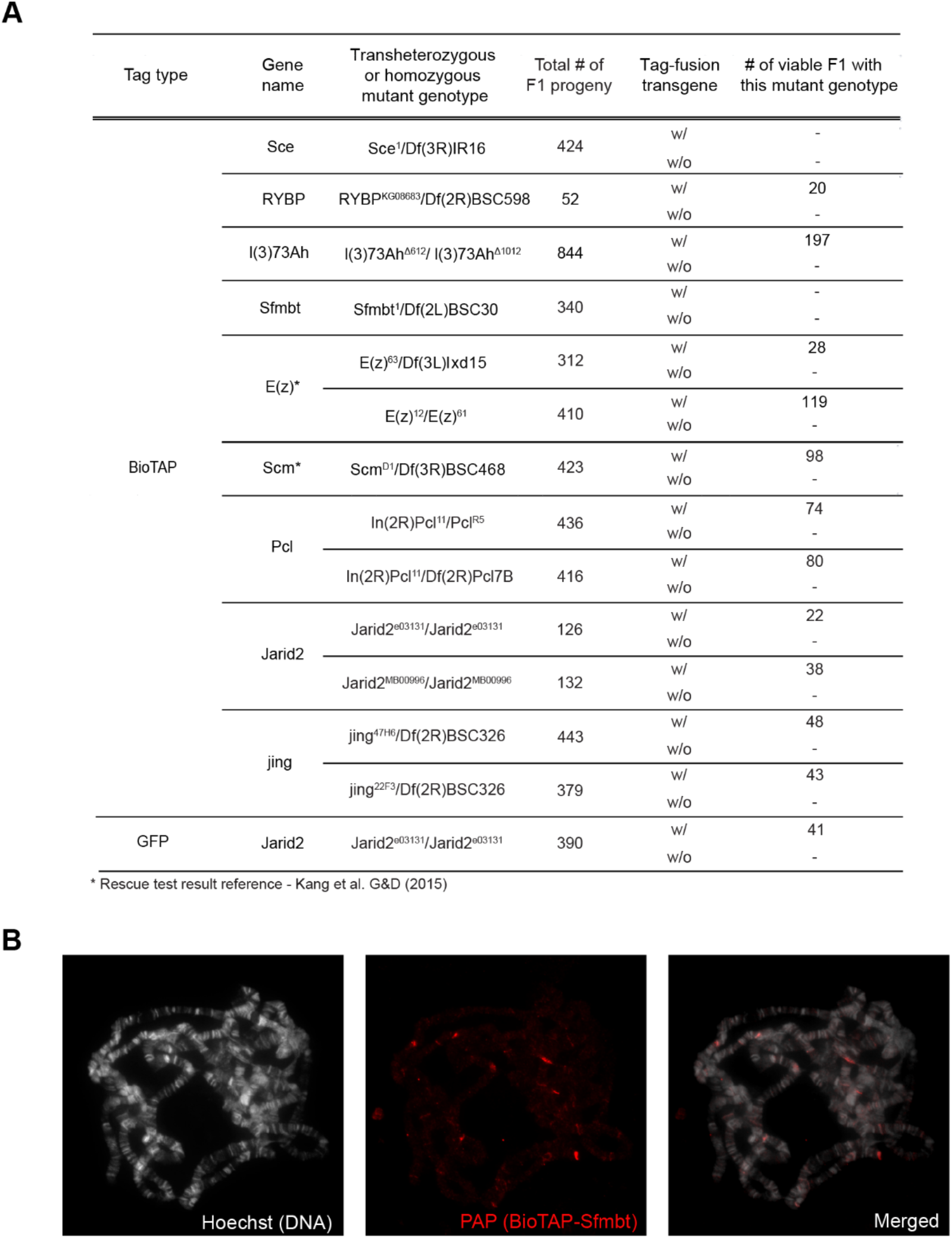
**(A)** Except for Kdm2, all mutants of proteins used in this study are lethal. Most BioTAP-tagged proteins rescued lethality of the mutants, but BioTAP-Sce (dRING) and BioTAP-Sfmbt failed in viability rescue tests. **(B)** Although it did not rescue Sfmbt mutant lethality, BioTAP-Sfmbt expressed in a wild-type background was detectable on larval polytene chromosomes, indicating the fusion protein could compete with its endogenous counterpart.

**Figure S3.**
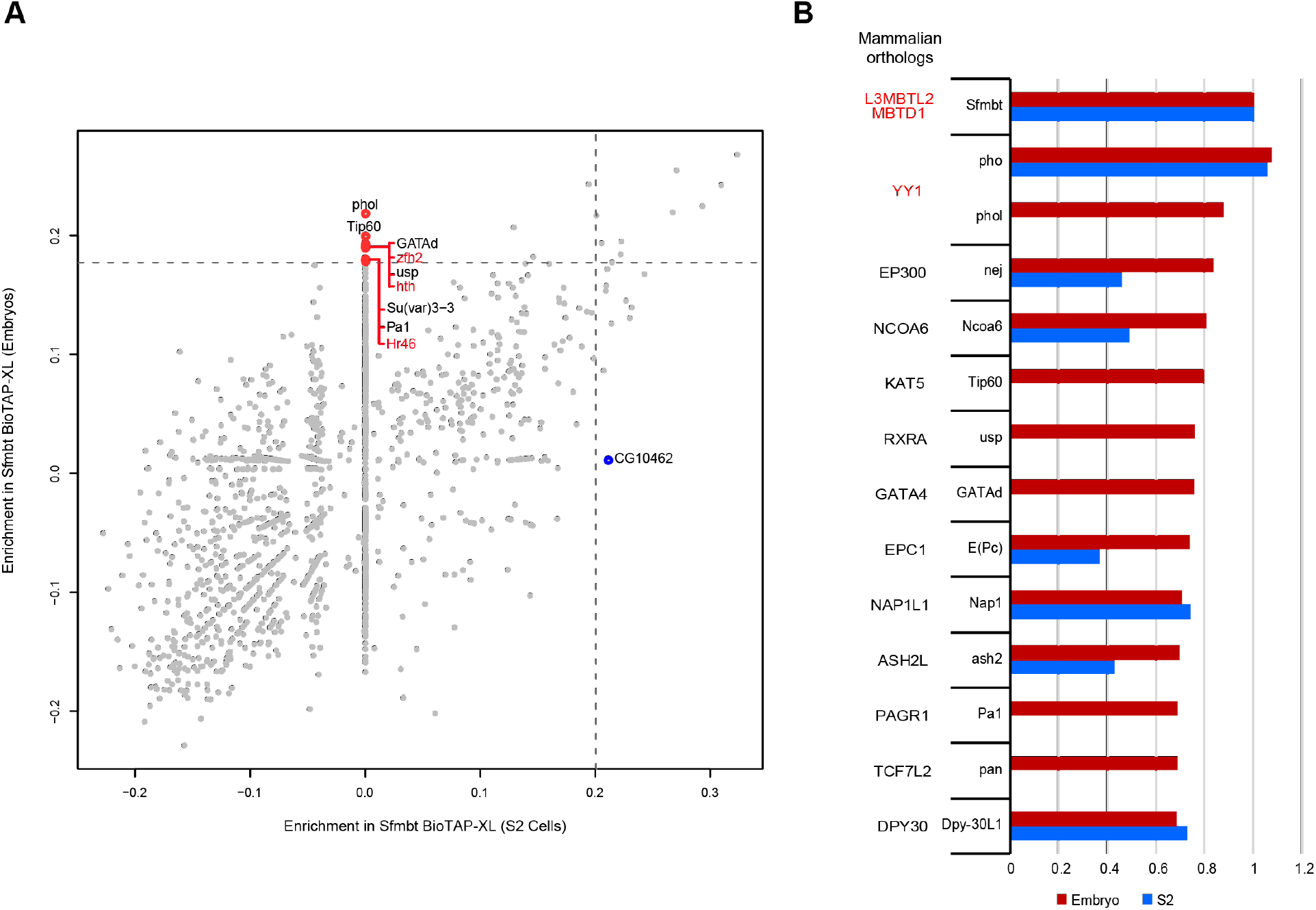
**(A)** Enrichment plot of Sfmbt-BioTAP-XL in S2 cells and embryos shows that there are embryo-specific interacting proteins. Proteins not expressed in S2R+ cells (zfh2, hth and Hr46) are highlighted in red. **(B)** The NSAF enrichment value of each highlighted protein is divided by the NSAF enrichment value of Sfmbt in embryos (red) and S2 cells (blue), and then graphed with the Sfmbt value set to 1.

**Figure S4.**
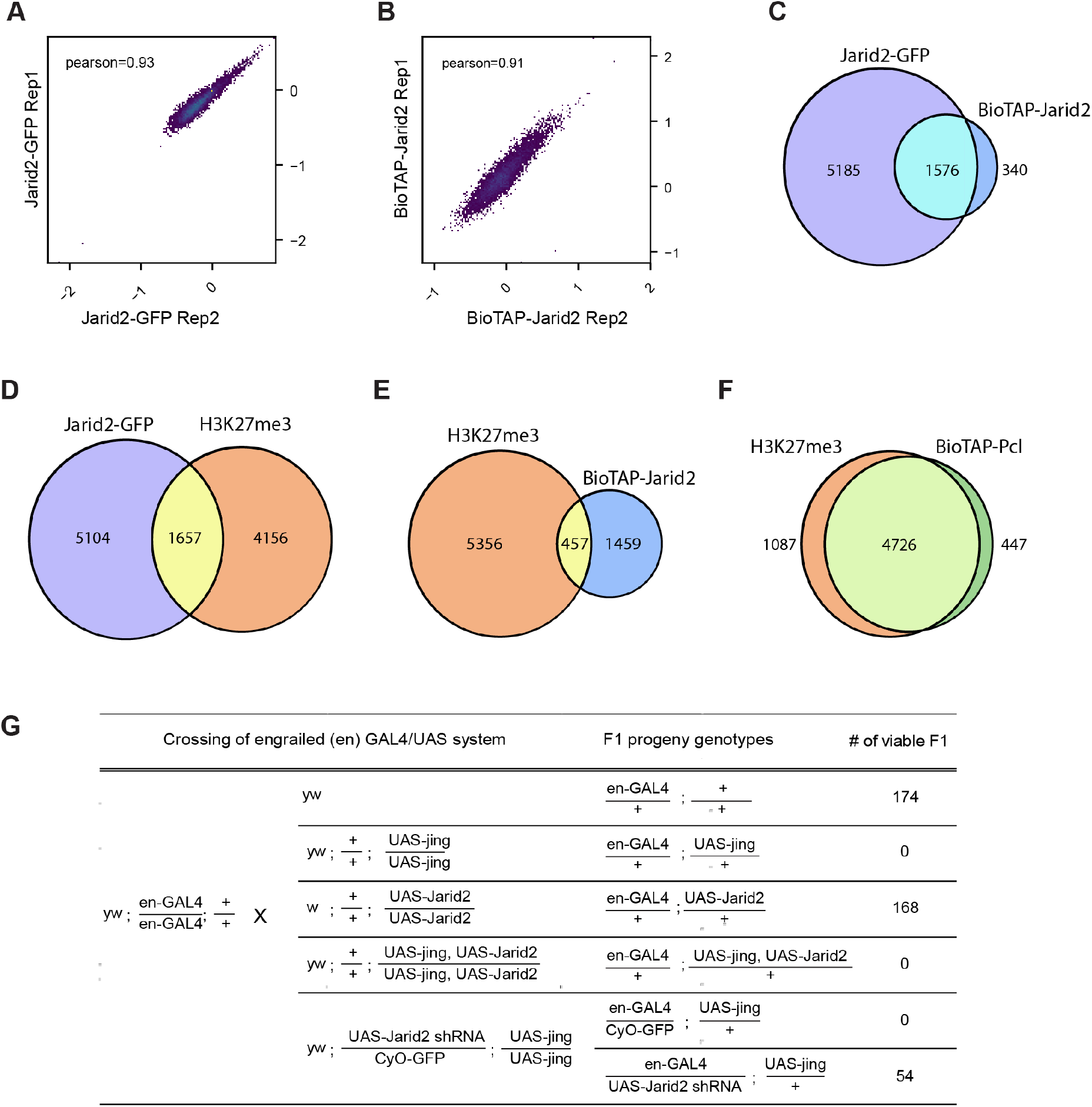
**(A-B)** Pearson correlation of ChIP-seq replicates (log2 ratio IP/Input) for Jarid2-GFP **(A)** and BioTAP-Jarid2 **(B)**. **(C-F)** Venn diagrams showing overlap of peaks between Jarid2-GFP and BioTAP-Jarid2 **(C)**, Jarid2-GFP and H3K27me3 **(D)**, H3K27me3 and BioTAP-Jarid2 **(E)**, and H3K27me3 and BioTAP-Pcl **(F)**. **(G)** Overexpression of jing using the GAL4/UAS system under control of *engrailed* regulatory sequences resulted in lethality at pupal stages. In F1 progeny from crosses in bold text, flies with wild-type wings (having both jing overexpression and Jarid2 knockdown) eclose, while adult flies with curly wings and GFP expression (having only jing overexpression) could not be found.

